# Desmosome-anchored intermediate filaments facilitate tension-sensitive RhoA signaling for epithelial homeostasis

**DOI:** 10.1101/2023.02.23.529786

**Authors:** Bageshri Naimish Nanavati, Ivar Noordstra, Suzie Verma, Kinga Duszyc, Kathleen J. Green, Alpha S. Yap

## Abstract

Epithelia are subject to diverse forms of mechanical stress during development and post-embryonic life. They possess multiple mechanisms to preserve tissue integrity against tensile forces, which characteristically involve specialized cell-cell adhesion junctions coupled to the cytoskeleton. Desmosomes connect to intermediate filaments (IF) via desmoplakin (DP) ^1,2^, while the E-cadherin complex links to the actomyosin cytoskeleton in adherens junctions (AJ) ^3^. These distinct adhesion-cytoskeleton systems support different strategies to preserve epithelial integrity, especially against tensile stress. IFs coupled to desmosomes can passively respond to tension by strain-stiffening ^4–10^, whereas for AJs a variety of mechanotransduction mechanisms associated with the E-cadherin apparatus itself ^11,12^, or proximate to the junctions ^13^, can modulate the activity of its associated actomyosin cytoskeleton by cell signaling. We now report a pathway where these systems collaborate for active tension-sensing and epithelial homeostasis. We found that DP was necessary for epithelia to activate RhoA at AJ on tensile stimulation, an effect that required its capacity to couple IF to desmosomes. DP exerted this effect by facilitating the association of Myosin VI with E-cadherin, the mechanosensor for the tension-sensitive RhoA pathway at AJ ^12^. This connection between the DP-IF system and AJ-based tension-sensing promoted epithelial resilience when contractile tension was increased. It further facilitated epithelial homeostasis by allowing apoptotic cells to be eliminated by apical extrusion. Thus, active responses to tensile stress in epithelial monolayers reflect an integrated response of the IF- and actomyosin-based cell-cell adhesion systems.

## Results

### Desmoplakin supports tension-activated RhoA signaling at adherens junctions

Although desmosomes and AJ make complementary contributions to mechanical homeostasis in epithelia ^5,7^, there is increasing evidence to suggest that these cytoskeletal-adhesion systems can interact functionally and biochemically ^14–16^. To test whether such interactions might affect the mechanical resilience of epithelia, we stimulated contractility in Caco-2 intestinal epithelial monolayers with the phosphatase inhibitor, calyculin A ^12,17,18^. Phase-contrast imaging revealed that control Caco-2 monolayers preserved cell-cell integrity for ~ 15-20 min, during which cell-cell contacts became more linear, consistent with increased tension (Fig 1A). Loss of epithelial integrity was first evident when sporadic points of separation between cells appeared. Separation then characteristically spread from these initiation points until clumps of cells had retracted from one another and rounded up.

**Figure 1.**
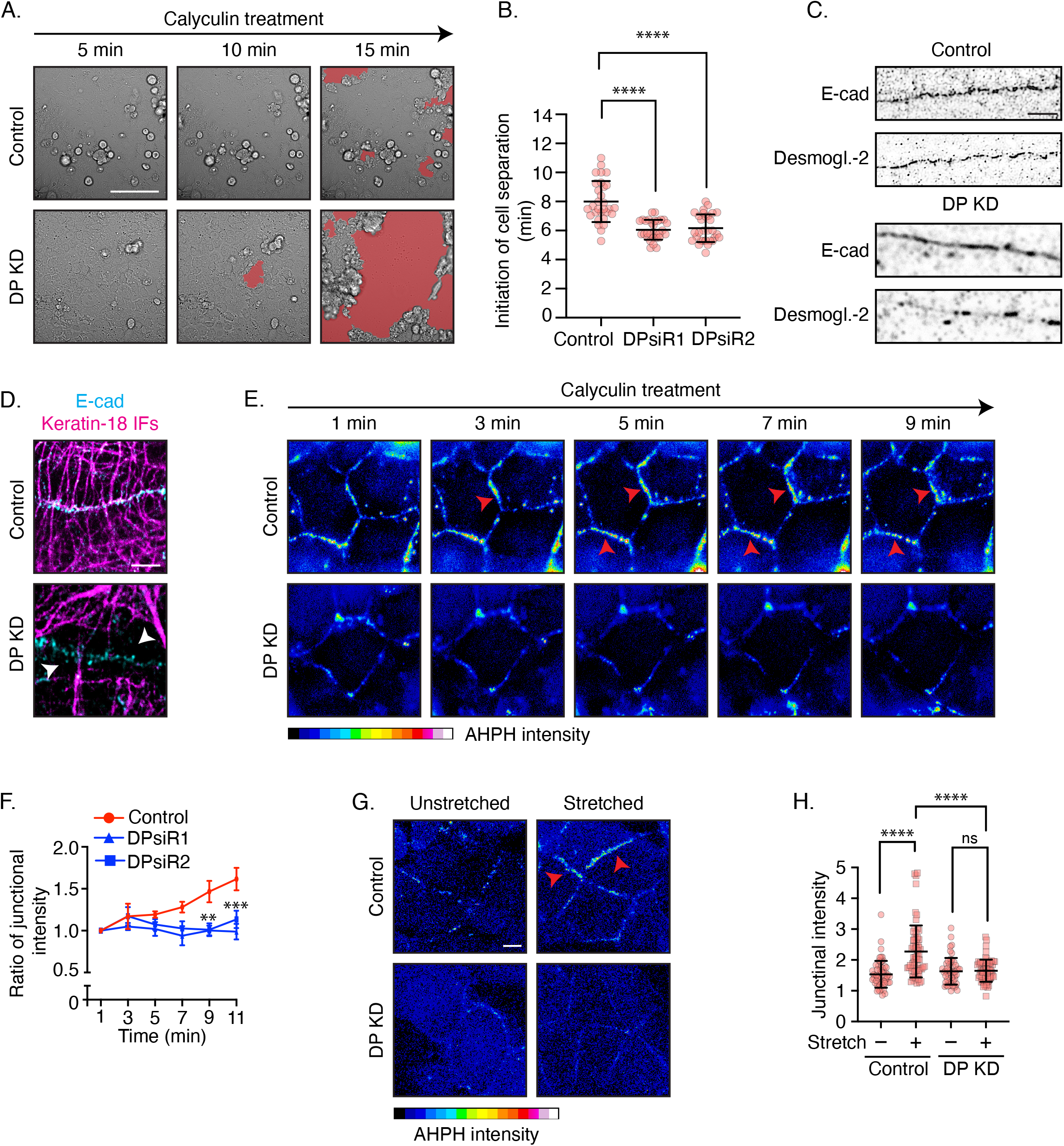
Desmoplakin supports tension-sensitive RhoA activation. (A,B) Response of control and DP-depleted (DPsiRNA 2) Caco2 monolayers to contractile stimulation with calyculin A (100 nM). A. Stills from phase-contrast movies; red arrows indicate the points of cell-cell separation. (B) Quantification of response to calyculin, measured as the time point when the first gaps appeared in the fields of view. (C,D) Effect of DP KD on cell junctions and IF anchorage. (C) Representative STED images showing distribution of E-cad and Dsg2 in control and DP-depleted monolayers. (D) Representative images of IFs and E-cad to visualise IF-anchorage in control and DP-depleted monolayers. E-cadherin was used to identify cell-cell contacts as Dsg2 was dramatically depleted by DP KD. (E,F) Response of active RhoA at cell-cell junctions to contractile stimulation with calyculin A (100 nM), detected with the AHPH location sensor for GTP-RhoA. (C) Representative images from movies; red arrows identify junctions where AHPH increases. (D) Quantification of junctional AHPH, normalized to junctional intensity at the beginning of each movie (0 min). (G,H) Response of junctional GTP-RhoA (AHPH) to monolayer stretch. Control and DP KD monolayers expressing AHPH were grown on flexible substrata before application of stretch. (E) Representative images and (F) quantification of change in junctional AHPH. Red arrows indicate junctions that showed an increase in AHPH. Error bars represent standard deviation with individual data points indicated from three independent experiments analysed using One-way ANOVA (B,F) or (D) Error bars represent standard error of mean calculated from mean of three independent experiments analysed using two-way ANOVA. ****p<0.0001, ***p<0.0005, **p<0.005, *p<0.05.

Then, we targeted DP ^19,20^ to disrupt the desmosome-IF system. We used two independent siRNAs that significantly reduced total cellular DP levels (knock-down, KD, Fig S1A). As expected ^1,2^, DP KD substantially reduced the number of desmosomes present at cell-cell contacts, compared with E-cadherin-containing AJ (Fig 1C), and disrupted the membrane-anchorage of IF (Fig 1D). The bulk of IF were found in the bodies of the DP KD cells and failed to extend into the cell-cell contacts (identified by E-cadherin), as was seen in controls. DP KD did not affect the integrity of monolayers under unstimulated steady-state conditions. However, loss of integrity was more evident when DP KD monolayers were stimulated with calyculin A (Fig 1A,B), where cell-cell separation began sooner than in controls. This was confirmed when we measured the time from addition of calyculin A to the first appearance of cell-cell defects (Fig 1B). Thus, DP helps Caco-2 monolayers resist tensile stresses, which is consistent with the generally-accepted role for the desmosome-IF system in supporting the mechanical resilience of epithelia ^2,5,21^.

However, tension-sensitive RhoA signaling at adherens junctions can also confer epithelial resistance to tensile stresses ^12^. Indeed, disabling this mechanoresponsive RhoA pathway accelerated the sensitivity of Caco-2 monolayers to calyculin A in a fashion that resembled the effect of DP KD. We therefore asked if DP KD might also affect tension-activated RhoA signaling at cell-cell junctions. We tested this using AHPH, a location biosensor for active, GTP-RhoA that is derived from the C-terminus of Anillin ^22,23^. The subcellular localisation and intensity of AHPH can be used as a read-out to measure localized changes in RhoA activity. We performed time-lapse imaging after calyculin A treatment to observe changes in GFP-AHPH (Fig 1E), focusing on the period before cell-cell separation became first evident. Quantification of fluorescence intensity revealed that the intensity of junctional AHPH increased continuously for ~15 min after calyculin A treatment in control cells (Fig 1F). However, the increase in junctional intensity was significantly compromised in DP KD cells, an effect that was confirmed with the two siRNAs (Fig 1E,F). Therefore, DP depletion hindered the junctional activation of RhoA in response to calyculin A treatment.

To reinforce this conclusion, we used an alternate method to apply tensile force by growing Caco-2 cell monolayers on a flexible substrate and then subjecting them to external stretch (10% sustained bi-axial, 10 min; Fig 1 G,H) ^12^. Junctional AHPH intensity increased significantly when control cells were stretched, indicating that application of external tension also activated RhoA in these cells (Fig 1G). However, junctional AHPH did not respond to stretch in DP KD monolayers (Fig 1 G,H). Together, these experiments indicate that DP is required for effective tension-activated RhoA signaling, providing an alternative mechanism for DP to confer resistance to tensile stress in epithelial monolayers.

### Desmosome-Intermediate filament linkage is necessary for tension to activate junctional RhoA

We then sought insight into how DP might support tension-activated RhoA signaling. Although DP is best-understood to mediate the physical linkage of IF to desmosomes, it can also interact with other cytoskeletal and signaling proteins ^24,25^. Therefore, we focused on testing whether IF coupling was critical for the impact of DP on tension-activated RhoA signaling.

We first adopted an independent loss-of-function approach by expressing DPNTP, a truncated mutant of DP, that has been reported to act as a dominant inhibitor that can uncouple IF from desmosomes ^21,26^. DPNTP contains the desmosome-binding N-terminus of DP but lacks the IF-binding domain. Transient expression of DPNTP in Caco-2 cells was confirmed by Western analysis (Fig S2A) and immunofluorescence demonstrated that the transgene localized to cell-cell junctions (Fig 2A, S2B). Spinning disc microscopy revealed that the subcellular localization of keratin IF was disrupted by DPNTP, with a clear gap being apparent between the bulk of IF in the bodies of the DPNTP cells and the cell-cell contacts (Fig 2A-C). Therefore, as previously reported ^21,26^, expression of DPNTP could uncouple IF from desmosomes.

**Figure 2.**
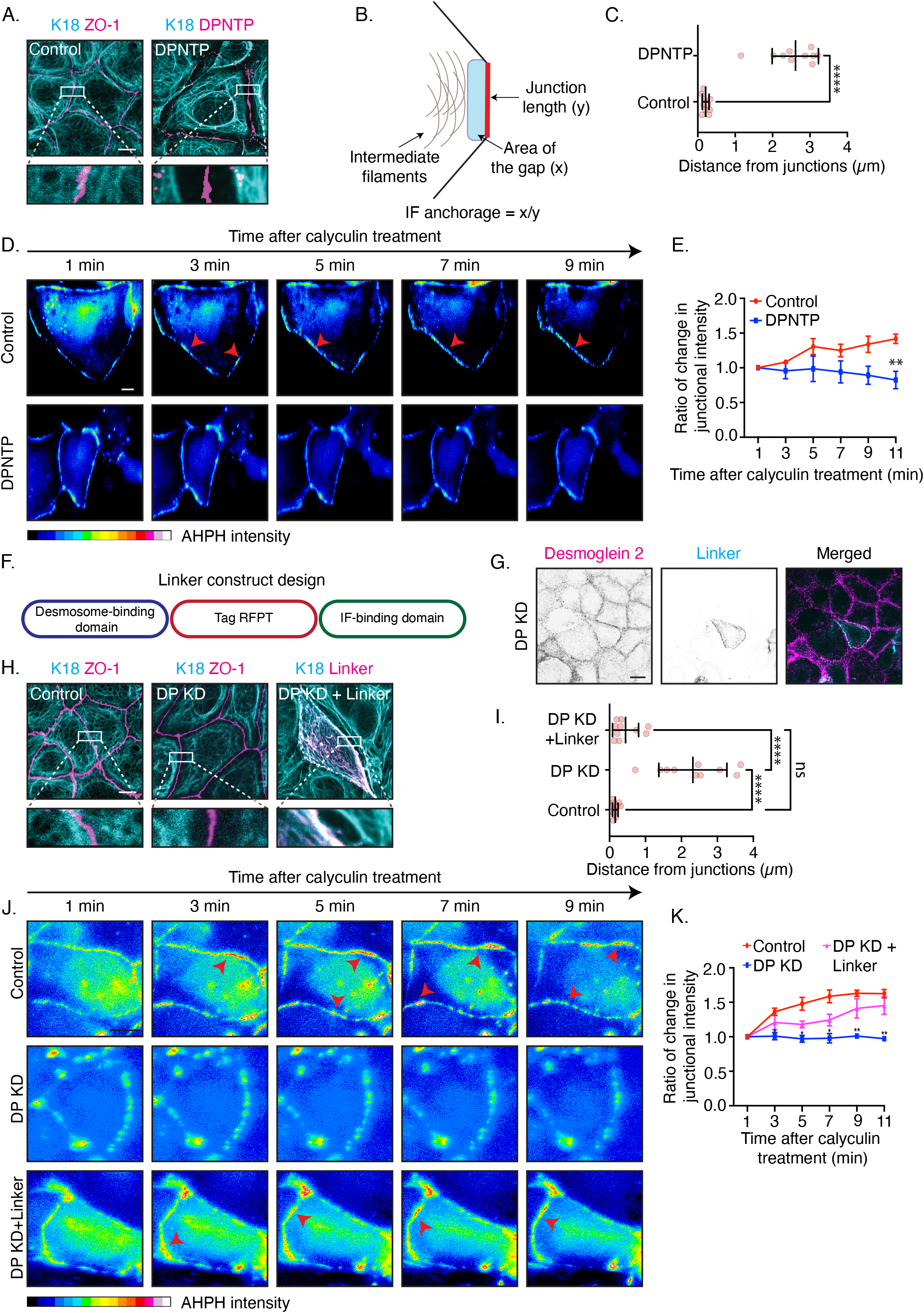
Linkage of intermediate filaments to desmosomes supports tension-activated RhoA. (A-C) Effect of DPNTP on intermediate filament (IF) to junction coupling. (A) Representative images of control and DPNTP-expressing cells immunostained for ZO-1 (used to mark cellcell contacts when desmosomes were perturbed) marking the cell-cell contacts and keratin 18. (B) Schematic of technique for quantifying IF anchorage to cell junctions. (C) Quantification of IF anchorage to cell junctions in control and DPNTP-expressing cells. (D,E) Effect of DPNTP on the junctional GTP-RhoA response to tensile stimulation. Control and DPNTP-expressing cells were transfected with AHPH and stimulated with calyculin (100 nM). (D) Representative images from time-lapse movies and (E) quantitation of junctional AHPH fluorescence (F,G) Development of the linker construct. Desmosome-binding domain is composed of 176 amino acids from the N-terminus of DP and the IF-binding domain is composed of 111 amino acids from the C-terminus of periplakin. (F) Schematic of the linker construct design (Linker-TagRFPT) and (G) its junctional localization in control and DP KD cells. (H,I) Effect of Linker-TagRFPT on IF anchorage to cell-cell junctions after DP KD. Control and DP KD cells expressing Linker-TagRFPT were immunostained for keratin 18 and ZO-1 to mark cell-cell contacts as desmosomes were perturbed. (H) Representative images and (I) quantification of IF-junction anchorage. (J,K) Stills of time-lapse imaging (J) and (K) quantification of junctional fluorescent intensity of AHPH after calyculin A treatment in control, DP KD and linker construct-expressing DP KD cells. Error bars represent standard deviation with individual data points indicated from three independent experiments analysed using unpaired t-test (C) or One-way ANOVA (I). (E,K) Error bars represent standard error of mean calculated from mean of three independent experiments analysed using two-way ANOVA. ****p<0.0001, ***p<0.0005, **p<0.005, *p<0.05.

Then we tested how DPNTP affected the RhoA response to tensile stress, measuring changes in junctional AHPH after stimulation with calyculin A. Whereas control cells showed a rapid and progressive increase in AHPH, this was abolished by DPNTP (Fig 2D,E). This suggested that linking desmosomes to IF might be important for the contribution of DP to tension-activated RhoA signaling. However, it remained possible that the demosomal-targetting action of DPNTP displaced other functions of full-length DP from the junctions.

Therefore, we developed a strategy to restore desmosome-IF coupling in DP KD cells. For this, we engineered a fusion protein consisting of the desmosomal-binding domain of DP and the keratin 8-binding domain of periplakin ^27^ (Linker-TagRFPT, Fig 2F, S2C,D), reasoning that this minimal construct could serve as an artificial linker to couple IF to desmosomes. Linker-TagRFPT expressed efficiently, as measured by Western analysis (Fig S2C) and localized to cell-cell junctions both in WT (Fig S2D) and DP KD cell lines (Fig 2H), as identified with its TagRFPT epitope tag. Importantly, Linker-TagRFPT effectively restored junctional coupling of IF to DP KD cells. The aberrant gap between cell-cell contacts and IF that was evident in DP KD cells was eliminated by expressing Linker-TagRFPT (Fig 2H, I). High-magnification views showed bundles of IF extending into the cell-cell contacts in the presence of Linker-TagRFPT, as were evident in controls but not KD cells (Fig 2H). Therefore, Linker-TagRFPT provided a tool to more selectively restore junctional coupling of IF to DP-depleted cells.

Importantly, Linker-TagRFPT, also restored the junctional RhoA response to tension when it was expressed in DP KD cells. Whereas DP KD abolished the junctional RhoA response to tension, KD cells expressing Linker-TagRFPT displayed a rapid-onset, progressive increase in AHPH (Fig 2J,K), similar to that seen in control cells. This indicated that coupling of IF to cell-cell junctions was a critical factor in allowing DP to support tension-activated RhoA signaling.

### DP supports the tension-sensitive RhoA mechanotransduction apparatus at AJ

Mechanical tension activates RhoA at adherens junctions via a mechanotransduction pathway where Myosin VI serves as a mechanosensor in association with E-cadherin ^12^. Application of tensile stress increases the interaction between Myosin VI and E-cadherin to initiate this signaling pathway. To test if DP might influence this mechanotransduction pathway, we first investigated its impact on the junctional recruitment of Myosin VI. Immunofluorescence showed that Myosin VI staining at cell-cell junctions increased when calyculin was applied to control monolayers (Fig 3A,B). However, DP KD monolayers showed reduced levels of Myosin VI at baseline, compared with controls, and these failed to increase further on stimulation with calyculin A. As the total cellular levels of Myosin VI were not altered by DP KD (Fig 3C), this suggested that DP influences the junctional recruitment of Myosin VI in response to tensile stress.

**Figure 3.**
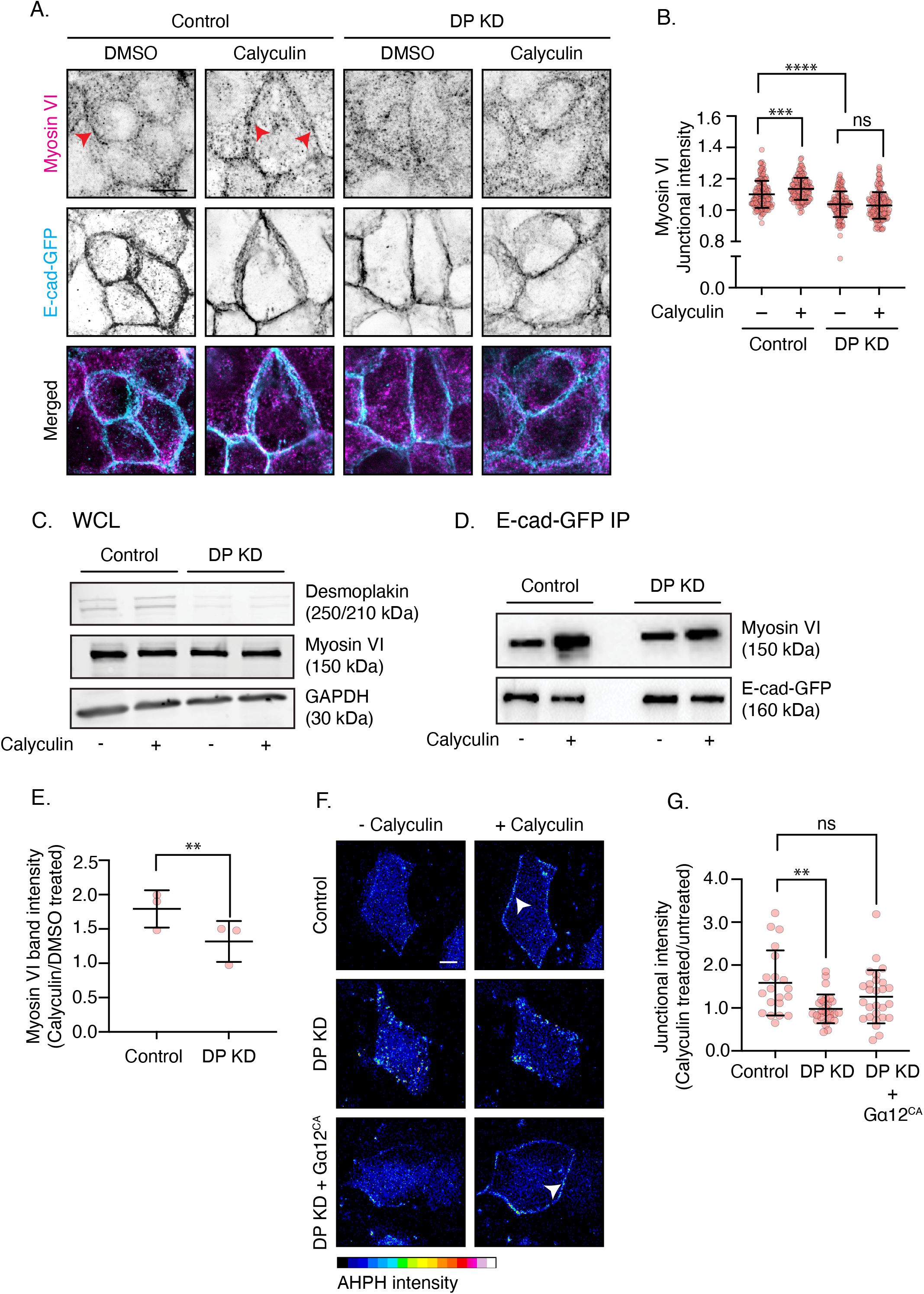
Desmoplakin facilitates the association between Myosin VI and E-cad for tension-sensitive RhoA signaling. (A,B) Effect of DP RNAi on Myosin VI recruitment to AJ following contractile stimulation. Control and DP KD cells were stimulated with calyculin (100 nM) and immunostained for Myosin VI and E-cadherin. (A) Representative images and (B) quantification of junctional fluorescence intensity. (C-E) Effect of DP on the tension-induced biochemical interaction between Myosin VI and E-cadherin. Experiments were performed in control and DP KD Caco-2 cells bearing CRISPR/Cas9-engineered E-cadherin-GFP. (C) Western analysis of whole cell lysates, (D) representative E-cadherin-GFP precipitates probed for Myosin VI and E-cadherin, and (E) quantitation of Myosin VI recruitment to E-cadherin in response to calyculin. (F,G) A constitutively-active Gα12 transgene (Gα12^CA^) rescues tension-activated RhoA signaling to DP KD cells. Junctional GTP-RhoA was identified with AHPH in control, DP KD and KD cells reconstituted with Gα12^CA^ (DP KD + Gα12^CA^). (F) Representative images of AHPH before and after stimulation with calyculin (100 nM; red arrows show increased AHPH at junctions) and quantitation of response (G). Error bars represent standard deviation with individual data points indicated from three independent experiments analysed using One-way ANOVA (B,G) or (E) Student t-test. ****p<0.0001, ***p<0.0005, **p<0.005, *p<0.05.

Then we tested if this reflected a change in the biochemical association of Myosin VI with E-cadherin (3D,E). For these studies, we used a Caco-2 cell line whose E-cadherin was engineered by CRISPR/Cas9 genome editing to bear a C-terminal GFP tag ^28^. E-cadherin-GFP was isolated with GFP-Trap and the protein complexes probed for Myosin VI. As previously reported, calyculin A increased the amount of Myosin VI that associates with E-cadherin ^12^. Application of tensile stress increases the interaction between Myosin VI and E-cadherin, but this was substantially compromised by DP KD (Fig 3D). To quantitate this, we normalized the Myosin VI signal to that of E-cadherin-GFP, then derived the ratio of the normalized Myosin VI signal in calyculin-treated cells compared with DMSO-treated controls. Recruitment of Myosin VI to E-cadherin increased by ~ 1.7 fold in control cultures that were stimulated by calyculin A and this was significant reduced in DP KD cells (Fig 3E). This indicated that DP supports the association of Myosin VI with E-cadherin in response to tensile stress.

These observations suggested that DP might influence RhoA through the E-cadherin-Myosin VI-based tension-sensing apparatus. If so, we reasoned that the impact of depleting DP would be reversed if we could activate downstream elements of the E-cadherin-Myosin VI pathway. To do this, we focused on the Gα12 G protein, which is recruited to the E-cadherin complex when tension is applied, and then engages the p114 RhoGEF to activate RhoA ^12^. We expressed a constitutively active form of Gα12 (Gα12^CA^) in DP KD cells and used the AHPH assay to measure the RhoA response to calyculin. Expression of Gα12^CA^ restored the tension-activated RhoA response in DP KD cells to control levels (Fig 3F,G), indicating that activation downstream in the E-cadherin-Myosin VI pathway could overcome the effects of DP depletion. Altogether, these results indicate that DP supports tension-activated RhoA signaling at AJ by facilitating the assembly of the E-cadherin and Myosin VI tension-sensing apparatus.

### DP supports neighbours of the dying cell to drive apoptotic extrusion

Together, these observations indicated that DP can link two processes that allow epithelia to respond to tensile stresses: coupling of IF to the membrane at desmosomes and tension-activated RhoA signaling at AJ. Both these processes have been recently implicated in the ability of epithelia to eliminate apoptotic cells by apical extrusion (apoptotic extrusion) ^17,29^. Therefore, we asked if the functional linkage for mechanotransduction that we have identified also contributed to apoptotic extrusion.

First, we confirmed that DP was necessary for apoptotic extrusion ^29^. We induced sporadic apoptosis by laser microirradiation of nuclei in E-cadherin-GFP Caco-2 monolayers. Time-lapse imaging confirmed that the apoptotic cell was apically expelled from control monolayers and the neighbours of the dying cell formed a rosette-like structure that sealed the monolayer (Fig 4A). However, as previously reported ^29^, apical extrusion failed in DP KD cultures, causing the apoptotic cells to be retained within the monolayers (Fig 4A,B). Then, we asked which cell population might require DP for apoptotic extrusion to occur. Effective extrusion involves active contributions from both the apoptotic cell and its neighbours ^30^. The apoptotic cell becomes hypercontractile when caspases activate pro-contractile kinases ^31–33^ to apply a tensile signal that activates mechanosensitive RhoA signaling in the neighbours ^17^. Therefore, we devised cell mixing experiments (Fig 4C) to test which of these cell populations required DP for apoptotic extrusion to occur.

**Figure 4.**
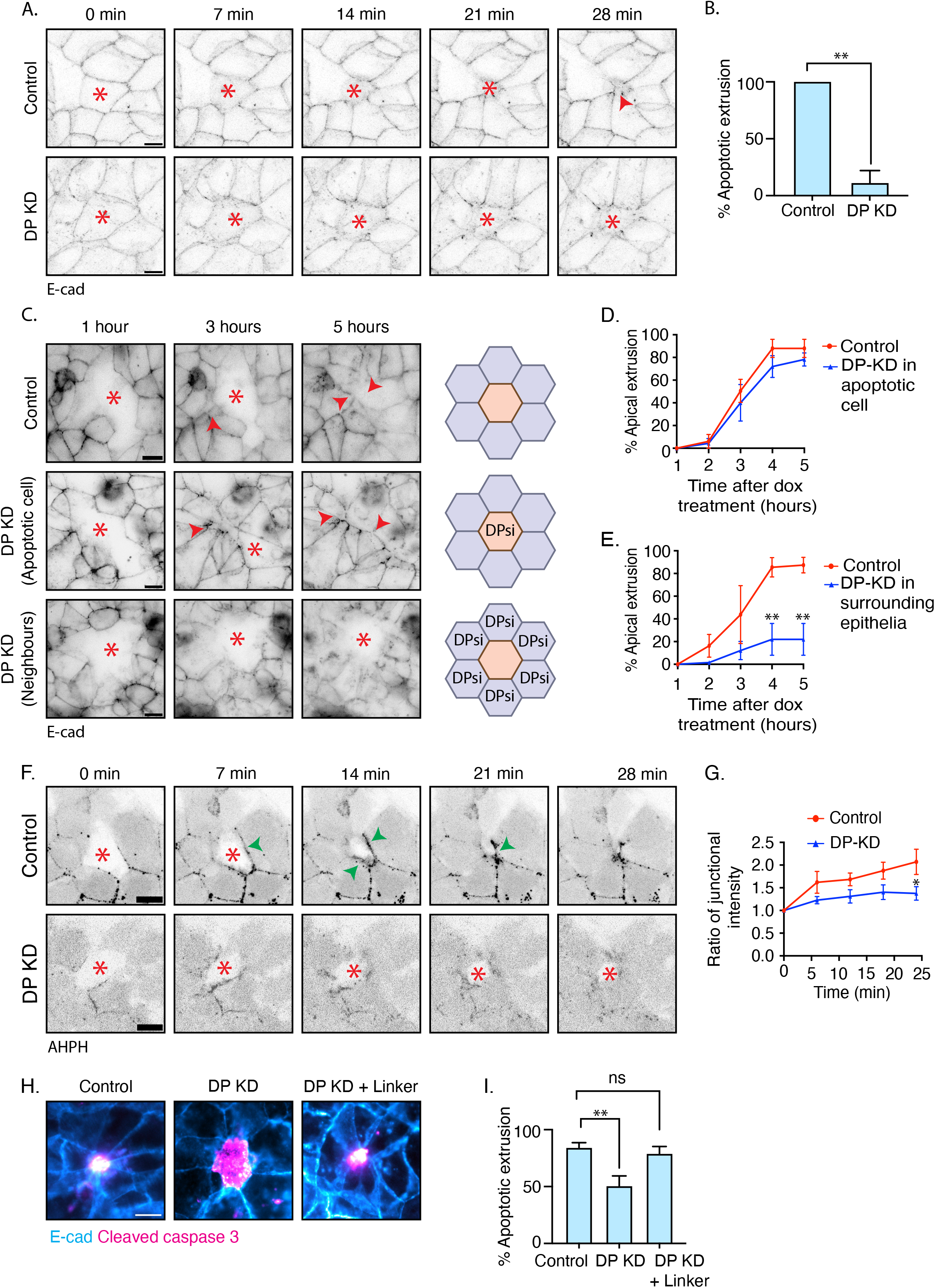
Desmoplakin supports RhoA activation in neighbours of apoptotic cells to drive their apical extrusion. (A,B) Effect of DP on the apical extrusion of apoptotic cells (apoptotic extrusion). Apoptosis was induced by laser micro-irradiation of selected cells (asterisks) in control and DP KD monolayers. (A) Stills from time-lapse movies and (B) quantitation of apoptotic extrusion. (C-E) DP is selectively required in epithelium surrounding apoptotic cells to support apical extrusion. Apoptosis was induced with Tet-inducible PUMA (see main text) and mixing experiments were used to deplete DP selectively from the PUMA cells (DP KD in PUMA cells, mixed 1:100 with WT E-Cad-GFP cells) or its surrounding epithelium (neighbours; PUMA cells with a WT-background mixed with DP KD in E-cadherin GFP cells, 1:100). (C) Stills from representative movies (imaged for 5 h after induction of PUMA with doxycycline and quantification of apical extrusion (% apoptotic cells that underwent extrusion) when DP was depleted in the PUMA (apoptotic cell; D) or in the surrounding epithelium (E). (F,G) Effect of DP on the neighbour-cell RhoA response during apoptotic extrusion. Apoptosis was induced by laser microirradation of AHPH-negative cells (asterisks) surrounded by AHPH-positive neighbours. (F) Stills from videos showing the RhoA response in neighbour cells, manifest as AHPH recruitment to the interface with the apoptotic cell (green arrowheads) and (G) Quantitation of the AHPH response at the apoptotic:neighbour interface. (H,I) Effect of rescuing IF anchorage on apoptotic extrusion in DP-depleted cells. Apoptosis was induced by 500nM/mL etoposide treatment for 6 hours in control, DP KD monolayers, and DP KD monolayers expressing Linker-TagRFPT. Cleaved caspase 3 (magenta) was used to mark apoptotic cells and E-cad-GFP (cyan) is used to mark cell boundaries. (H) Representative images of cells undergoing apoptosis (I) Quantitation of apical extrusion (% apoptotic cells undergoing apical extrusion). Red asterisk: cells undergoing apoptosis, Red arrow: formation of rosette-like structure by neighbours of the dying cell (C), Green arrow: changes in junctional fluorescent intensity Error bars represent standard error of mean from three independent experiments analysed using One-way ANOVA (B,I) or analysed using two-way ANOVA (D,E,G). ****p<0.0001, ***p<0.0005, **p<0.005, *p<0.05.

For these experiments, apoptosis was induced in Caco-2 cells that express the pro-apoptotic protein, p53 Upregulated Modulator of Apoptosis (PUMA-mCherry) under control of the Tet-ON promoter system ^17^. PUMA^TetON^ cells were then mixed with E-cadherin-GFP cells at a ratio (1:100) that created small groups of PUMA^TetON^ cells surrounded by PUMA^TetON^-null cells. Induction of PUMA-mCherry expression by addition of doxycycline (5 hr) caused the sporadic PUMA^TetON^ cells to undergo apoptosis and be apically extruded from the monolayer (Fig 4C). We then counted the proportion of apoptotic cells that underwent extrusion as an index of apoptotic extrusion.

To test if effective extrusion required DP in the apoptotic cell, we transfected PUMA^TetON^ cells separately with DP siRNA and mixed these with E-cadherin-GFP cells that had not been transfected with siRNA. Surprisingly, DP-depleted PUMA^TetON^ cells were extruded as efficiently as PUMA^TetON^ cells that retained DP (Fig 4C,D). This implied that DP was not required in the apoptotic cell for extrusion to occur. Then, to test a role for DP in the neighbour cell population, we mixed PUMA^TetON^ cells with E-cadherin-GFP cells that had been separately transfected with DP siRNA. Here there was a significant decrease in the proportion of apoptotic cells that were extruded (Fig 4C,E). This indicated that DP is selectively required in the epithelia around apoptotic cells for apical extrusion to be effectively achieved.

Mechanosensitive RhoA signaling is activated in the neighbours of apoptotic cells to mediate extrusion ^17,34^. Therefore, we hypothesized that its impact on RhoA signaling might explain why DP was required for the neighbours to mediate extrusion. Accordingly, we used AHPH to test if DP siRNA affect the RhoA response in neighbour cells (Fig 4F,G). To ensure that we only measured GTP-RhoA in the neighbours, we mixed WT cells with GFP-AHPH expressing cells (1:100 ratio), then induced apoptosis by laser microirradiation of cells not expressing GFP-AHPH. In control monolayers, AHPH in neighbours progressively accumulated at their immediate interface with the apoptotic cell as extrusion occurred (Fig 4F,G). However, this RhoA response was significantly compromised by DP KD. Therefore, the capacity for DP to support tension-activated RhoA signaling provided a potential mechanism for it to facilitate apoptotic extrusion.

Finally, since coupling of IF was necessary for DP to support tension-activated RhoA signaling, we tested if this also participated in apoptotic extrusion. For this, we induced apoptosis (etoposide, 6 hr) in DP KD cells that expressed Linker-TagRFPT and compared the proportion of apoptotic cells that were extruded with that in controls or DP KD cells (Fig 4H, I). Strikingly, Linker-TagRFPT restored the extrusion that was impaired by DP KD to control levels. Thus, junction-IF coupling was necessary for apoptotic extrusion, as it was for tension-activated RhoA signaling.

## Conclusion

Desmosomes and AJs play key, often complementary ^5,7^, roles in allowing tissue integrity to be preserved against tensile stresses. Our current findings reveal that the desmosome-IF system can influence epithelial homeostasis through cross-talk with mechanotransduction at AJ. Thus, IF coupling by DP supports the tension-activated RhoA signaling pathway that assembles at E-cadherin-based adhesions to reinforce the junctional actin cytoskeleton ^12^. This impact of DP on AJ mechanotransduction was functionally significant in our studies, as it enhanced monolayer resistance to tensile stresses and supported the homeostatic process of apical extrusion. Importantly, in these cases the contribution of DP appeared to reflect its ability to couple IF to cell-cell junctions, as it was restored with our linker construct. Future studies will be necessary to elucidate the detailed mechanism that is responsible for this connection between two specialized adhesion-cytoskeletal systems. Potentially, the DP-IF linkage could influence the mechanical loading that is sensed by the E-cadherin-Myosin VI pathway. This may be through indirect cross-talk between desmosomes and AJ, although binding to plakoglobin ^35^ may allow DP to be also recruited to AJ. More generally, our findings potentially identify a new way for the passive mechanical properties of IF to modulate AJ signaling. This reinforces our growing awareness of the diverse mechanisms that can link the desmosome-IF and AJ-actomyosin systems systems ^14–16,20^. IF and F-actin networks can be structurally linked by proteins, such as spectroplakins, while IF-desmosomal linkages influence cell mechanics driven by actomyosin and interact with actin regulators ^8,36,37^. Conversely, cortical tension can influence desmosomal biogenesis ^16^. Tension-activated RhoA signaling contributes to morphogenetic processes that range from apoptotic extrusion ^17^ to collective epithelial cell migration ^38^. Therefore, the pathway that we have identified reveals a new avenue for the IF cytoskeleton to contribute to epithelial homeostasis.

## Acknowledgements

We thank our colleagues for their support and advice throughout this project. Our work was supported by grants from the National Health and Medical Research Council of Australia (1136592 and 1163462), The Australian Research Council (DP220103951) and the National Institutes of Health (R01AR043380; R01AR041836; R01CA228196 to KJG). I.N. was supported by the European Molecular Biology Organization (EMBO ALTF 251-2018). Microscopy was performed at the ACRF/IMB Cancer Research Imaging Facility created with the generous support of the Australian Cancer Research Foundation.

## Author contributions

Conceptualization, B.N.N., I.N., K.D., K.J.G., A.S.Y; Methodology, B.N.N., I.N., K.D.; Formal analysis, B.N.N.; Investigation, B.N.N., S.V., I.N.; Writing, B.N.N., K.J.G., A.S.Y.; Visualization, B.N.N., I.N.; Supervision, I.N., A.S.Y.; Funding acquisition, K.J.G., A.S.Y.

## Supplementary Figure Legends

**Figure S1, related to Figure 1.**

(A) Optimisation of DP knockdown. Western blot analysis to visualise expression of DP at two different time points after transfection with control siRNA or siRNAs targeting DP.

**Figure S2, related to Figure 2**

(A,B) Expression of DPNTP in Caco-2 monolayers. Cells were transiently transfected with DPNTP-GFP 24 hours before processing for Western analysis (A) or immunofluorescence (B): a representative image of DPNTP-expressing cell co-stained with Dsg2. Scale bar = 10μm.

(C,D) Expression of Linker construct in control and DP siRNA cells. (E) Western blot analysis confirming the expression of the linker construct at the predicted molecular weight of 60kDA in control and DP KD cells. (F) Representative image of the linker construct expressed in wild-type Caco-2 cells and - co-stained with Dsg2 to confirm the junctional localisation of the linker construct. Scale bar = 10μm

## Methods

### RESOURCE AVAILABILITY

#### Lead contact

Further information and requests for resources and reagents should be directed to and will be fulfilled by the lead contacts, Alpha Yap

#### Materials availability

All reagents generated in this study will be made available upon request to the lead contact.

#### Data availability

All data generated in this study will be made available upon request to the lead contact.

### EXPERIMENTAL MODEL AND SUBJECT DETAILS

#### Cell culture and transfection

Human colorectal adenocarcinoma (Caco-2) cells were obtained from ATCC (HTB-37) and cultured in Roswell Park Memorial Institute (RPMI) media supplemented with 10% Fetal Bovine Serum (FBS), 1% L-Glutamine, 1% MEM non-essential amino acids and 100 units/mL of penicillin/streptomycin. Human Embryonic Kidney 293 (HEK293T) cells were cultured in Dulbecco’s Modified Eagle Medium (DMEM) supplemented with 10% FBS, 1% L-Glutamine, 1% MEM non-essential amino acids and 100 units/mL of penicillin/streptomycin. All cells were grown in a humidified chamber at 37°C in 5% CO_2_ atmosphere.

Caco-2 cells were transfected with plasmids using Lipofectamine 3000 (Thermo Fisher Scientific #L3000015) according to the manufacturer’s protocol. For siRNA transfection, cells were transfected with Lipofectamine RNAiMAX (Thermo Fisher Scientific #13778150) according to the manufacturer’s protocol. When transfected with plasmids, cells were analysed 24 hours post transfection. When transfected with siRNA, cells were analysed 48-72 hours post transfection.

#### Stable cell lines and lentiviral transduction

Caco-2 E-cad-GFP knock-in cell line was established using CRISPR-Cas9 technology as described previously ^1^. PUMA^TET-ON^ Caco-2 cells, and GFP-AHPH Caco-2 cells were generated as described previously ^2^.

Caco-2-Linker-TagRFPT cell line was generated using lentiviral-based transduction technique. To generate lentiviruses, HEK-293T cells were transfected with a lentiviral expression vector, pTripZ and packaging constructs, pMDLg/pRRE, RSV-Rev and pMD.G using Lipofactamine 2000 according to the manufacturer’s protocol. The virus was concentrated from the supernatant of the HEK-293T cells after 48 hours of incubation, using concentrating spin columns. Subsequently, Caco-2 cells were infected with the concentrated virus. TagRFPT-positive Caco-2 cells were selected using FACS Aria Cell Sorter (Queensland Brain Institute, the University of Queensland).

### METHOD DETAILS

#### DNA constructs

To synthesize pC1 Linker construct, a gblock was synthesized containing 176 AA from the N-terminus of DP and 111 AA from the C-terminus of periplakin. The gblock was cloned into p-C1 vector by restriction digestion with Age1 and BamH1. Linker construct was subsequently cloned into a pLVX-IRES-Puro lentiviral backbone by restriction digestion with Age1 and Hpa1.

#### CalyculinA treatment

Intrinsic contractility was increased by treating 90-100% confluent Caco-2 cell monolayers with 100nM/mL calyculinA (Abcam #ab141784) for 10-15 min at 37°C.

#### Induction of apoptosis

##### a. Laser injury

E-cad-GFP Caco2 cells were cultured on 35 mm glass-bottom dish. When cells reached 90-100% confluency, apoptosis was induced by nuclear irradiation using a Mai Tai two-photon laser. To record cellular responses to apoptosis, time-lapse imaging was performed continuously for 30 min. AnnexinV (Life Technologies #A23204) was used as a marker to identify cells undergoing apoptosis.

##### b. Tet-inducible PUMA

PUMA^TetON^ cells were mixed with E-cad-GFP-Caco2 cells at a 1:100 ratio and cultured on 35 mm glass bottom dishes, to create small clusters of PUMA cells surrounded by E-cad-GFP-Caco2 cells. When cells reached 90-100% confluency, apoptosis was induced in PUMA^TetON^ cells by treating the mixed monolayer with 1μg/mL doxycycline. To allow sufficient time for expression of PUMA to induce apoptosis and record cellular responses to apoptosis, timelapse imaging was performed for 5 hours after the treatment with doxycycline.

##### c. Etoposide treatment

E-cad-GFP Caco-2 cells or E-cad-GFP Linker-TagRFPT Caco2 cells were cultured on coverslips in 6-well plates. When the cells reached 90-100% confluency, they were treated with 500nM/mL etoposide (Adooq Biosciences LLC #A10373) for 6 hours at 37°C. After the treatment, cells were fixed with 100% methanol and stained for Cleaved Caspase 3 to mark apoptotic cells and GFP to stain for E-cad to identify cell borders.

#### Immunofluorescence

Caco-2 cells were fixed using 4% Paraformaldehyde (PFA) in cytoskeleton stabilisation buffer (10mM PIPES at pH 6.8, 100mM KCl, 300mM sucrose, 2mM EGTA and 2mM MgCl_2_) for 20 min at room temperature (RT) or were fixed using 100% methanol for 10 min at −20°C. Staining for Myosin VI, DP, keratin-8, keratin-18, Dsg2, tagRFPT and GFP required pre-permeabilisation of cells (0.1% Triton in PBS) for 3 min on ice before fixation with PFA. Subsequently, the fixed cells were blocked in 3% Bovine Serum Albumin (BSA) in PBS at RT for 1 hour. Following this, cells were incubated with primary antibody for 1 hour at RT. For Myosin VI and Dsg2 primary antibodies, the cells were incubated overnight at 4°C. Then, the cells were washed with 0.05% Tween in PBS before incubating with the corresponding secondary antibodies at RT for 1 hour. Following this, the coverslips were mounted using Prolong gold (Genesearch #8916BC) with or without DAPI.

#### Immunoprecipitation

For GFP-trap experiments, E-cad-GFP Caco-2 cells were cultured on 10 cm dishes. After 72-96 hours of plating, the cells were treated with calyculinA or DMSO for 10 min at 37°C.

Then, the cells were lysed for 30 min in lysis buffer (1% NP40, 10mM Tris-HCl pH7.4, 150mM NaCl, 2mM CaCl_2_, 1x protease inhibitor). Meanwhile, GFP beads (Chromotek #gta-20) were blocked in 3% BSA in Tris-Buffered Saline (TBS) for 4 hours at 4°C. The cell lysates were incubated with GFP beads overnight at 4°C. After washing the beads with lysis buffer several times, the protein complexes were resolved using SDS-PAGE.

#### Western blotting

Caco-2 cells were lysed in 1x lysis buffer (4x: 200mN Tris-HCl, 40% glycerol, 8% SDS and 0.4% Bromophenol Blue) and incubated at 95°C for 9 min. The cell lysates were resolved in 8% or 12% SDS polyacrylamide gels, then transferred to nitrocellulose membrane and blocked with 5% milk in TBS for 1 hour at RT. The membranes were incubated with primary antibodies overnight at 4°C. Later, the membranes were incubated with the corresponding horseradish (HRP) – conjugated secondary antibodies for 1 hour at RT. The blots were resolved using Supersignal West Pico PLUS Chemiluminescent Substrate (Thermo Fisher #34579) and imaged on Biorad Chemidoc. For Myosin VI the membranes were incubated with Alexa Fluor conjugated secondary antibodies. These blots were imaged using Odyssey CLX.

#### Microscopy

Fixed samples were imaged on an Andor Dragonfly inverted confocal spinning disc microscope operated using Fusion and Imaris software. Images were acquired using 60x, 1.4 NA Plan Apo objective. Fixed samples were imaged on Upright Zeiss Axiolmager widefield microscope operated using Zeiss ZEN blue 3 software. Images were acquired using 40x, 0.95 NA Plan Apo objective or 63x, 1.40 NA Plan Apo objective.

Time-lapse imaging after calyculinA treatment and for PUMA-induced apoptotic extrusion was performed on Nikon Ti-E deconvolution microscope operated using NIS elements software was used. Images were acquired using 20X, 0.75 NA Plan Apo objective. Time-lapse imaging for laser-induced apoptotic extrusion was performed on Zeiss LSM 710 Meta confocal microscope equipped with Mai Tai multi-photon laser operated using Zen Black software. Images were acquired using 43X, 1.3 NA Plan Apo objective.

Subcellular distribution of E-cadherin and Desmoglein 2 was visualised on Leica SP8 STED 3X FLIM super resolution microscope operated using Leica LASX software. Images were acquired using HC Plan Apochromat 93×1.30 glycerol objective.

### QUANTIFICATION AND STATISTICAL ANALYSIS

#### Image analysis

##### a. Initiation of cell separation

The initiation of cell separation was quantified by noting the time point after calyculinA treatment when there was formation of gap in the cell monolayers.

##### b. Intermediate filaments anchorage

The anchorage of IFs was measured as the average distance between IFs and cell junctions. The average distance was determined by dividing the area of the gap between IFs and cell junctions to the length of the same junction.

##### c. Apoptotic extrusion

Apoptotic extrusion was classified successful when: (1) the apoptotic marker-positive cell was expelled out of the monolayer, and (2) the neighbours of the apoptotic cell formed a rosette-like structure to avoid formation of gaps in the monolayer. Apoptotic extrusion was classified as a failure when (1) the apoptoticmarker-positive cell was retained in the monolayer, and (2) the neighbours of the apoptotic cell failed to elongate to form a rosette-like structure leading to formation of gaps in the monolayer.

##### d. Junctional intensity

Junctional intensity of AHPH and Myosin VI was measured by taking a ratio of the junctional intensity and cytoplasmic intensity. For measuring the changes in junctional intensity of AHPH after calyculinA treatment or induction of apoptosis, we calculated the ratio of the junctional intensity of AHPH after stimulation (with calyculin A or induction of apoptosis) compared with its junctional intensity before stimulation.

#### Statistical analysis

All the experiments have been performed independently at least 3 times. Statistical parameters for the individual experiments are included in the specific figure legends. The parameters include sample size, statistical test performed and statistical significance.

