## Supplementary figures and images for "Desmosome-anchored intermediate filaments facilitate tension-sensitive RhoA signaling for epithelial homeostasis"

### Supplementary Figures

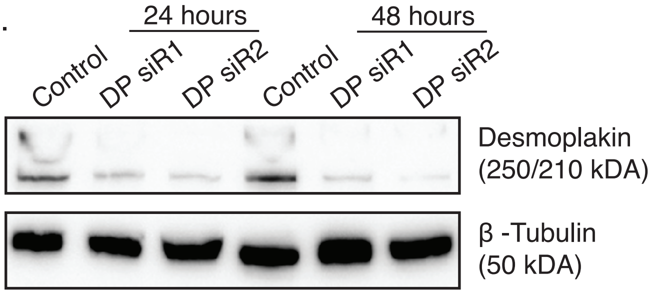

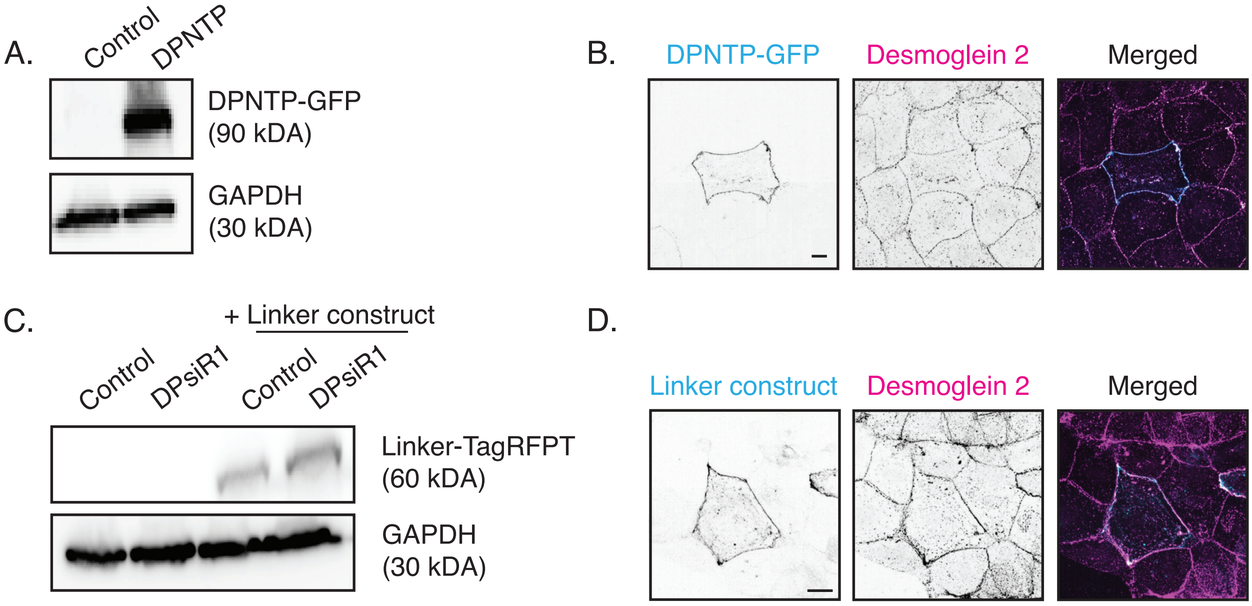
